# Functional analysis of evolutionary human methylated regions in schizophrenia patients

**DOI:** 10.1101/540294

**Authors:** Niladri Banerjee, Tatiana Polushina, Anne-Kristin Stavrum, Vidar Martin Steen, Stephanie Le Hellard

## Abstract

**Background:** Recent studies have implicated variations in DNA methylation in the aetiology of schizophrenia. Genome-wide scans in both brain and blood report differential methylated regions (DMRs) and positions (DMPs) between patients with schizophrenia and healthy controls. Previously, we reported that DMRs where human specific methylation (hDMR) has occurred over evolutionary time are enriched for schizophrenia-associated markers (SCZ_hDMR). However, it is unknown whether these human specific DMRs show variable methylation in patients with schizophrenia.

**Methods:** Using publicly available data, we investigate if human specific DMRs that harbour genetic variants associated with schizophrenia are differentially methylated between cases and controls.

**Results:** We find statistically significant (*p* < 1e-4) methylation difference in schizophrenia associated human specific DMRs (SCZ hDMR) between brain samples of cases and controls. However, we fail to find evidence of similar differences in methylation in blood samples.

**Conclusion:** Regions that are evolutionarily important for human species and that are associated with schizophrenia, also show difference in methylation variation in the brain in patients with schizophrenia.

## Introduction

Schizophrenia is a devastating psychiatric illness that has a global lifetime prevalence of 1% [1] and affects patients in the prime of their youth. It has no cure and treatment relies on antipsychotic drugs that can cause several side effects such as weight gain [2], obesity [3] and metabolic syndrome [4]. These side effects are believed to induce higher mortality amongst patients with schizophrenia with average life expectancy reducing by nearly 20 years [5,6].

Much research has been undertaken in the last decade to unravel the genetics of the disease in the hope of finding better treatment options for the patients. Research has revealed the disease to have complex aetiology with the latest genome-wide association study (GWAS) reporting over a hundred loci genome-wide [7]. While these results have brought more understanding about the regions involved in the disease, much work remains to elucidate the biological mechanisms. A majority of the genome-wide hits are in the non-coding regions [7] and efforts are on to investigate the role of epigenetic factors in these regions [8]. For instance, differences in DNA methylation pattern amongst patients with schizophrenia, compared to healthy controls at both candidate gene loci [9–12] and at the global genome-wide level [13–17] have been reported.

DNA methylation has also been investigated in studies of epigenetic inheritance [18–20] and paleo-epigenetics [21–23]. For example, Gokhman and colleagues [23] looked at methylation differences between humans and our extinct hominid cousins-Neanderthals and Denisovans. Although most of the methylated regions were similar between the three hominids, 2000 regions were differentially methylated amongst the three hominid species. Building up on their study, we found that the human specific differentially methylated regions (hDMRs) show an enrichment of genetic variants associated with schizophrenia (compared to the rest of the genome [24]). While regions differentially methylated between human and great apes [25] do not show such an enrichment.

Other genetic studies have also suggested that genomic markers of recent evolution are enriched for schizophrenia associated single nucleotide polymorphisms (SNPs) [28–30]. Thus together with our results this has narrowed the evolutionary hypothesis of the schizophrenia proposed by TJ Crow and others [26,27] to a possible effect of recent evolution in the maintenance of schizophrenia in humans.

We have now investigated whether these human specific DMRs which are in regions of genetic association with for schizophrenia could also show differences in methylation between patients with schizophrenia and controls.

## Materials & Methods

### Evolutionary Differentially Methylated Region (DMR) data

Gokhman et al. [23] published a list of DMRs they term as human-specific differentially methylated regions (hDMRs). They identified these DMRs by comparing human methylome against Neanderthal and Denisovan methylomes. They represent human lineage specific methylation markers since the divergence of the common ancestor of Neanderthals and Denisovans.

### Evolutionary DMR selection for analysis

We have previously reported the enrichment of association of the hDMRs for schizophrenia [24]. Among the 891 hDMRs, we focus on DMRs containing SNPs significantly associated with schizophrenia. We first determine an experiment-wide significance threshold to select the SNPs and DMRs. About 27,000 SNPs were located in or in linkage-disequilibrium (LD) with h_DMRs [24]. We followed the procedure of Moskvina et al. [31] and estimated 2048 independent SNPs that with a Bonferroni correction at α = 0.05 gives an experiment-wide threshold of 2.46×10e-5. Nine DMRs were found to harbour SNPs that passed this threshold and were carried forward in the analysis - *DMR10, DMR127, DMR203, DMR204, DMR236, DMR237, DMR291, DMR526, DMR527* **(Additional File 1, 2)** To account for the sparse and irregular coverage of array probes in these DMRs, we included the ∼3kbp flanking regions of the 9 DMRs, which gave a final list of 65 probes that covered a maximum these 9 DMRs **(Additional File 1).**

### Brain and Blood Methylation Data

Data for methylation variation in brain were obtained from publicly available datasets at GEO accession numbers : GSE61107 [14], GSE6143 [16], GSE61380 [16], GSE74193 [17] and for blood at GSE80417 [15]. Normalized beta matrix was used for all datasets except for GSE61107 where raw IDAT files were available.

### Data Processing

All the publicly available methylation data analysed in this study were obtained from the Illumina HumanMethylation450 BeadChip platform. IDAT files were available for the study by Wockner and colleagues [14]. We removed probes that failed at a detection *p*-value of 0.05 in 50% of the samples. Subsequently, functional normalization [32] was performed using the *minfi* [33] package in R [34]. During pre-processing we found that 4 DMRs had only a single overlapping probe. **(Additional File 1)**. These probes are also in the list from Zhou et al. [35] where they recommend removing the probes as they contain SNPs that could interfere with probe performance **(Additional File 1).** We chose not to remove these probes, as doing so would eliminate the only methylation signal within the DMRs. Details of the other DMRs containing probes suggested for masking are available in additional file 1. Similarly, we did not need to remove probes on the sex chromosomes since none of the evolutionary DMRs selected were on these chromosomes.

The study by Pidsley et al. [16] comprised of two different brain datasets obtained from the Douglas Bell Canada Brain Bank (DBCBB) and London Neurodegenerative Diseases Brain Bank (LNDBB). These were analysed separately similar to a comparative analyses performed by Wockner and colleagues [13].

The study by Jaffe et al. [17] initially comprised of more than 650 samples in total. After removing duplicate and pre-natal data, we focused on a final set of 191 schizophrenia patient samples with 231 controls with a similar age distribution **(Supplementary Table T1)**.

Normalized data from Pidsley et al., 2014 [16] and Hannon et al., 2016 [15] were used directly.

### Visualization of DMRs

DMRs were visualized using the *Gviz* package [36] in R [34]. Genomic coordinates for SNPs and RefSeq genes in hg19 were obtained from the UCSC Table Browser [37]. CpG island information was obtained using the Annotation Hub package [38]. Probe annotation for the Illumina HumanMethylation450 BeadChip was obtained from study by Zhou et al. [35].

### Significance testing of DMPs and DMRs

We implemented pipeline using *limma* [39,40] package in R [34] with empirical Bayes [41] to determine the differentially methylated positions (DMPs) that showed statistically significant differences in methylation between patients with schizophrenia and controls. Details of the linear models implemented for each of the datasets may be found in supplementary file.

To determine whether the specific evolutionary DMRs selected showed statistically significant differences in methylation at the region level, we implemented *comb-p* [42], which combines adjacent *p*-values and performs false discovery rate adjustment. This makes it ideal for analysing data from DNA methylation arrays where probes are irregularly spaced. Furthermore, *comb*-p allows one to test custom defined regions for statistical significance. In our case, we implemented *comb-p* for the individual DMRs. We report here the *p*-values from one-step Sidak multiple testing correction performed using the *region_p* programme.

### Epigenetic Annotation

The 18-state chromatin annotation was obtained from the Roadmap Epigenome project [43] for sample E073 that contains a mixture of dorso-lateral prefrontal cortex tissues from two people aged 75 and 81. Additionally, chromatin state annotation was obtained from peripheral blood mononuclear cells from sample E062. The BED file containing the genome coordinates of the 18 state model was intersected with the coordinates of the 9 DMRs in R [34] using the *Genomic Ranges* package [44].

## Results

### Evolutionary enriched DMRs show methylation variation in patients with schizophrenia

We utilised the brain methylation datasets from studies conducted by Wockner et al.. [14] (N = 48, 24 cases with schizophrenia, 24 controls), Pidsley et al.. (LNDBB: N = 46, 22 cases with schizophrenia and 24 controls; DBCBB: N = 33, 18 cases with schizophrenia and 15 controls) [16] and Jaffe et al. (N = 322, 191 cases with schizophrenia and 231 controls) [17]. All three studies utilized the prefrontal cortex for methylation analyses. We find evidence of variable methylation in the SCZ_hDMRs in all three datasets.

In the largest brain methylation dataset from Jaffe et al. [17], we find methylation differences between cases and controls for *DMR203* (*p* = 7.29e-17) **(Figure 1)**, *DMR291* (*p* = 1.756e-4) and *DMR526* (*p* = 1.019e-4) **(Table 1)**. This dataset might be more reliable due to all the factors that could be included in the regression model which is not the case for other datasets (**Supplementary Methods).** *DMR203* also shows different level of methylation between cases and controls in the DBCBB dataset from Pidsley et al. **(Supplementary Figure S1, Table 1)**. *DMR127* shows differences in methylation in the brain datasets from Wockner **(Figure 2)** and Pidsley **(Supplementary Figure S2, S3),** but the direction of effect is opposite in the two samples. Furthermore, although *DMR127* replicates in both the LNDBB and DBCBB datasets from Pidsley, it survives multiple testing correction only in the LNDBB dataset **(Table 1)**

**Figure 1.**
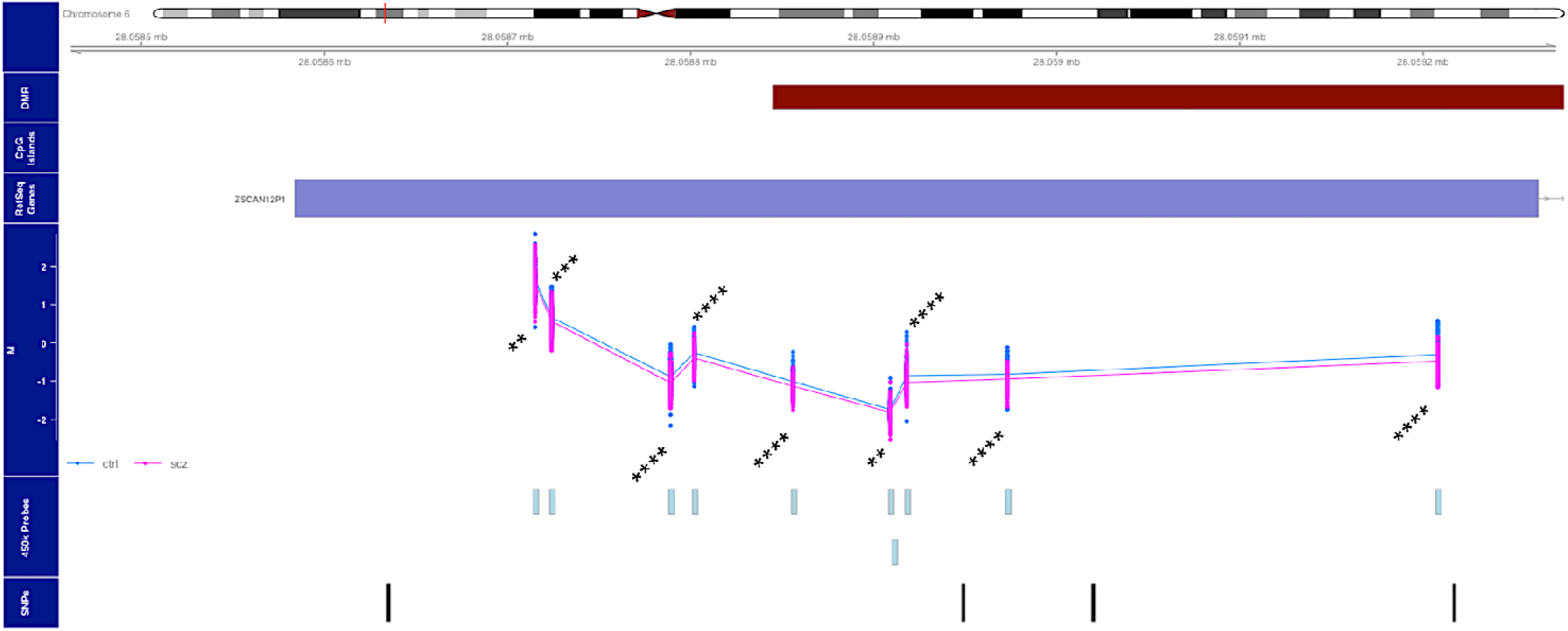
Methylation variation in *DMR203,* in samples from Jaffe et al. Figure depicts methylation variation in patients with schizophrenia in the evolutionary *DMR203* (maroon, *p* =) in the prefrontal cortex. Schizophrenia methylation (pink) compared with controls (blue) from Jaffe et al. [17]. Significant differentially methylated positions (DMPs) depicted ** (*p <*0.005), *** (*p <*0.0005), and **** (*p <*0.00001)

**Table 1.** *Comb-p* results across all evolutionary DMRs and datasets. Table depicts the *p*-values from Sidak multiple correction for all the DMRs across all the brain methylation datasets obtained from *comb-p*. Statistically significant DMRs are denoted with * (*p <* 5e-2), ** (*p <* 1e-3) and *** (*p <* 1e-4). *DMR526* and *DMR527* lacked probes, possibly removed during pre-processing by Pidsley et al. [16] and as such could not be tested.

|  |  |  | (LNDBB) | (DBCBB) |  |
| --- | --- | --- | --- | --- | --- |
| **DMR10** | 1:2378997-2379771 | 1 | 1 | 1 | 1 |
| **DMR127** | 3:10805466-10809820 | 1.824e-2* | 1.078e-2* | 0.5602 | 1 |
| **DMR203** | 6:28058845-28060341 | 1 | 1 | 5.064e-3* | 7.29e-17*** |
| **DMR204** | 6:28833654-28834570 | 0.997 | 1 | 1 | 1 |
| **DMR236** | 7:1952455-1953065 | 1 | 1 | 1 | 1 |
| **DMR237** | 7:2048069-2049043 | 1 | 1 | 1 | 1.367e-3* |
| **DMR291** | 8:38287269-38289920 | 0.9733 | 1 | 1 | 1.756e-4** |
| **DMR526** | 14:103745950-103746209 | 1 | - | - | 1.019e-4** |
| **DMR527** | 14:104006646-104008448 | 1 | - | - | 1 |

**Figure 2.**
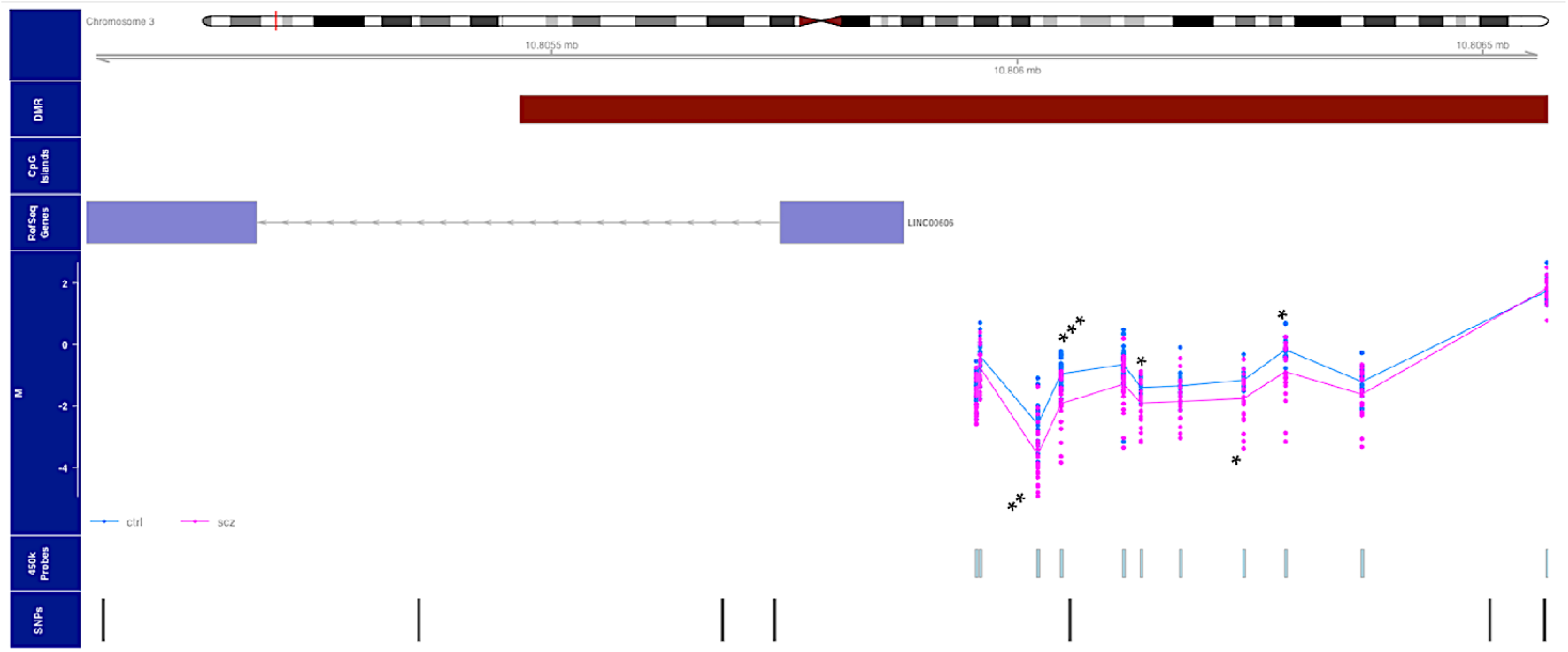
Methylation variation in *DMR127,* in samples from Wockner et al. Figure depicts methylation variation in patients with schizophrenia in the evolutionary DMR (maroon) in the prefrontal cortex. Schizophrenia methylation (pink) compared with controls (blue) from Wockner et al. [14]. Significant differentially methylated positions (DMPs) depicted as * (*p <*0.05), ** (*p <*0.005) and *** (*p <*0.0005)

### Methylation variation does not appear in blood samples

We performed the same analyses in blood samples obtained from Hannon et al. [15] (N = 675, 353 cases with schizophrenia and 322 controls). We found no evidence of different methylation in any of the SCZ_hDMRs in blood samples **(Supplementary Figures S4, S5, Supplementary Table T2)**. However, our analysis was limited due to lack of information on all the covariates from the Hannon study [15].

### Evolutionary enriched DMRs occur in diverse genome annotations

We annotated the evolutionary enriched DMRs using data from the Roadmap Epigenomics Consortium [43]. We utilised the 18-state chromatin model from sample E073 [17] that comprises of data from a mixture of the dorsolateral prefrontal cortex from two persons aged 75 and 81 years old. The annotation suggests *DMR127* to be present in an active enhancer that overlaps *LINC0606; DMR203* overlaps an active transcription start site for *ZSCAN12P1* **(Supplementary Table T3)**. Additionally, we find that *DMR291* overlaps weak enhancers in *FGFR1* **(Supplementary Table T3)**. In addition, using Hi-C data from the hippocampus [45] in the 3D Genome Browser [46] we observed that *DMR526* is potentially located in a regulatory region for the gene *MARK3* **(Supplementary Figure S6)**. We also annotate the DMRs using the chromatin state data in blood from E062. We find that the DMRs that are significantly differentially methylated in the brain datasets occur in heterochromatin regions in blood **(Supplementary Table T3).**

## Discussion & Conclusion

We report here the investigation of DNA methylation differences between cases with schizophrenia and controls in regions that show differential methylation between modern humans and Neanderthal/Denisovan and which are associated with schizophrenia. We observe the difference in methylation for several of these regions in brain samples but not in blood samples.

*DMR203* shows statistically significant methylation difference in patients with schizophrenia compared to controls that survives multiple testing correction using the Sidak procedure in *comb-p*. We find evidence in the largest set of brain samples from Jaffe et al. [17] (*p* < 1e-16, Sidak correction) and the DBCBB samples *(p*< 0.05, Sidak correction) from Pidsley et al. [16]. However, the direction of effect is not consistent. This could be due to lack of power in the two samples (type I error in both), however it could also be due to different cell composition between the two studies. We observe *hypo*-methylation in samples from Jaffe et al. [17] **(Figure 1)** and *hyper*-methylation in the DBCBB samples [16] **(Supplementary Figure S1)**. The *DMR203* is within the extended MHC region that is known to harbour the genetic variants with the strongest association with schizophrenia. This DMR overlaps with the pseudo gene *ZSCAN12P1.* The gene has also been implicated in schizophrenia and autism spectrum disorder in a recent meta-analysis [47].

*DMR127* shows statistically significant differences in methylation in both the Wockner dataset [14] and the LNDBB dataset from Pidsley et al.[16], but the direction of effect is opposite between the two studies. We observe *hypo*-methylation in the schizophrenia samples from Wockner [14] **(Figure 2)** while the LNDBB samples from Pidsley et al.[16] show *hyper*-methylation in the patient samples **(Supplementary Figure S2).** This could again be due to lack of power in the two samples or as reported previously, because of different cell composition between the two studies [13]. Interestingly, *DMR127* also shows significantly differentially methylated probes (DMPs) in the DBCBB dataset from Pidsley et al. **(Supplementary Figure S3)**, but the region does not survive corrections for multiple testing multiple testing in *comb-p* **(Table 1)**. The region of *DMR127* covers the long intergenic non-protein coding RNA 606, *LINC0606*. Since there is different methylation in patients with schizophrenia in this region, it is possible that the regulation of this gene is important in the aetiology of schizophrenia. The gene is actively expressed in the brain and occurs upstream of *SLC6A11* and downstream of *ATP2B2,* both of which are also highly expressed in the brain and have been implicated in various neurological disorders such as epilepsy [48], autism [49] and schizophrenia [50]. However, *DMR127* does not replicate in the largest brain methylation dataset from Jaffe et al. [17]. It is possible that due to increased sample size and power in the dataset from Jaffe et al. [17], the *hypo*- and *hyper*-methylation observed in the smaller datasets of Wockner et al. [14] and the LNDBB dataset [16] get averaged out. There may also be specific sub-populations of patients with schizophrenia who show either *hypo* or *hyper*-methylation in this region. The other possibility is the lack of identical cell types analysed across all the brain datasets. This is a challenge for the field of psychiatric genetics that the brain is a heterogeneous tissue filled with varying levels of neurons, glial cells and astrocytes. This makes comparative epigenetics studies difficult to execute and implement.

Finally we find some methylation differences in three additional DMRs – *DMR237 (p <* 1e-2, Sidak correction), *DMR291 (p <* 1e-3, Sidak correction) and *DMR526 (p <* 1e-3, Sidak correction) in the dataset from Jaffe et al. [17]. *DMR237* occurs in the intronic region of *MAD1L1,* but the significance appears to be driven by a single DMP, instead of a broad change across the DMR (data available on request). *DMR291* overlaps the gene *FGFR1* while *DMR526* is in a non-coding region of the genome. *FGFR1* has been previously implicated in schizophrenia [51,52]. *DMR526* does not overlap with any genes but may be involved in long-range regulation of *MARK3* determined through Hi-C genome-wide contact matrix **(Supplementary Figure S6)**,

An important caveat of the present analysis is that many of the DMRs have low probe coverage **(Additional File 1)**. For instance, *DMR526* which shows some difference in methylation in the dataset from Jaffe et al. is represented by only one probe. Several other DMRs had limited probe coverage which is a major limitation of our analyses. In the normalized datasets from Pidsley et al. [16], the only probe in *DMR526* was missing. This also implies a subsequent limitation of the statistical test in *comb-p*. For multiple testing correction using the Sidak procedure, the algorithm relies on probe coverage in the DMRs. Thus, filtering out probes may have an impact on the final *p*-values reported by Sidak correction.

When looking at the blood samples [15] from patients with schizophrenia, we fail to find evidence of any methylation variation in these evolutionary DMRs. This is unlikely due to lack of power as the sample size from Hannon et al. [15] is comparable to that of Jaffe et al. [17]. The lack of replication in blood might instead suggest that the disease is more specific to the brain and thus we may not expect to observe difference in blood, at least in the particular SCZ_hDMRs analysed in the present study. This may be supported by the fact that the 18-state chromatin annotation from REMC for peripheral blood mononuclear cells shows the most significant SCZ_hDMRs to be in quiescent and repressed chromatin regions **(Supplementary Table T3)**. This is supported by data from GTEx [53] that shows higher expression of the genes in the specific SCZ_hDMRs in brain over blood **(Supplementary Figure S4, S5)**. Furthermore, we cannot rule out that there may be other h_DMRs that do not contain significant SCZ SNPs, but may show disrupted methylation variation in blood. Additionally we cannot rule out that the methylation variation observed in brain samples is due to anti-psychotic medications [54]. An additional point to consider is that the variation we observe in the evolutionary DMRs in brain may in fact be driven by methylation quantitative trait loci (mQTL). Such mQTLs may act in *cis* upto 1Mb from the CpG site [55]. Previous research into foetal brain mQTLs has demonstrated a significant enrichment with schizophrenia associated regions [56]. Future research should look into the influence of genotypes affecting methylation variation in evolutionary DMRs.

In conclusion, we observe that regions of the human genome whose methylation is specific to human evolution and are enriched for schizophrenia associated markers show disrupted methylation in brain samples of patients with schizophrenia. This disrupted methylation is not observed in blood samples. Our results suggest dysregulation of methylation in novel regions of the genome in patients with schizophrenia in brain tissue. Future research should be carried out with higher probe density in the evolutionary DMR regions or use whole genome methylation sequencing technologies.

