## Supplementary data, methods and figures for "Functional analysis of evolutionary human methylated regions in schizophrenia patients"

**SUPPLEMENTARY MATERIAL**

a. NORMENT, Department of Clinical Science, University of Bergen, Bergen, Norway

b. Dr. Einar Martens Research Group for Biological Psychiatry, Department of Medical Genetics, Haukeland University Hospital, Bergen, Norway

**Correspondence to:** Prof. Stéphanie Le Hellard, Department of Clinical Medicine, Laboratory Building, Haukeland University Hospital, N-5021 Bergen, Norway.

**CONTENTS**

**Supplementary Methods	3**

**Supplementary Figures S1-S9	4**

FIGURE S1	4

FIGURE S2	6

FIGURE S3	8

FIGURE S4	10

FIGURE S5	10

FIGURE S6	11

FIGURE S7	12

**Supplementary Tables T1-T3	13**

TABLE S1	13

TABLE S2	13

TABLE S3	14

### Supplementary Methods

#### *Regression Models*

### 1. Wockner et al

### *M value ~ age +* *gender + post-mortem interval + status*

### 2. Pidsley et al (DBCBB + LNDBB)

### *M value ~ age + gender + status*

### 3. Hannon et al

### *M value ~ age + gender + status*

Here it is important to acknowledge the fact we did not have all the covariate information used by Hannon et al. in the original study for each of the samples. The original study also included covariates of smoking and cell composition.

### 4. Jaffe et al

### *M value ~ age + gender + status + race + embryonic stem cell proportion + neural progenitor cell proportion + dopaminergic cell proportion + neuronal cell proportion + non neuronal cell proportion*

Detailed covariate information for each of the patient and controls in this dataset was publicly available and was used in the model.

### Supplementary Figures

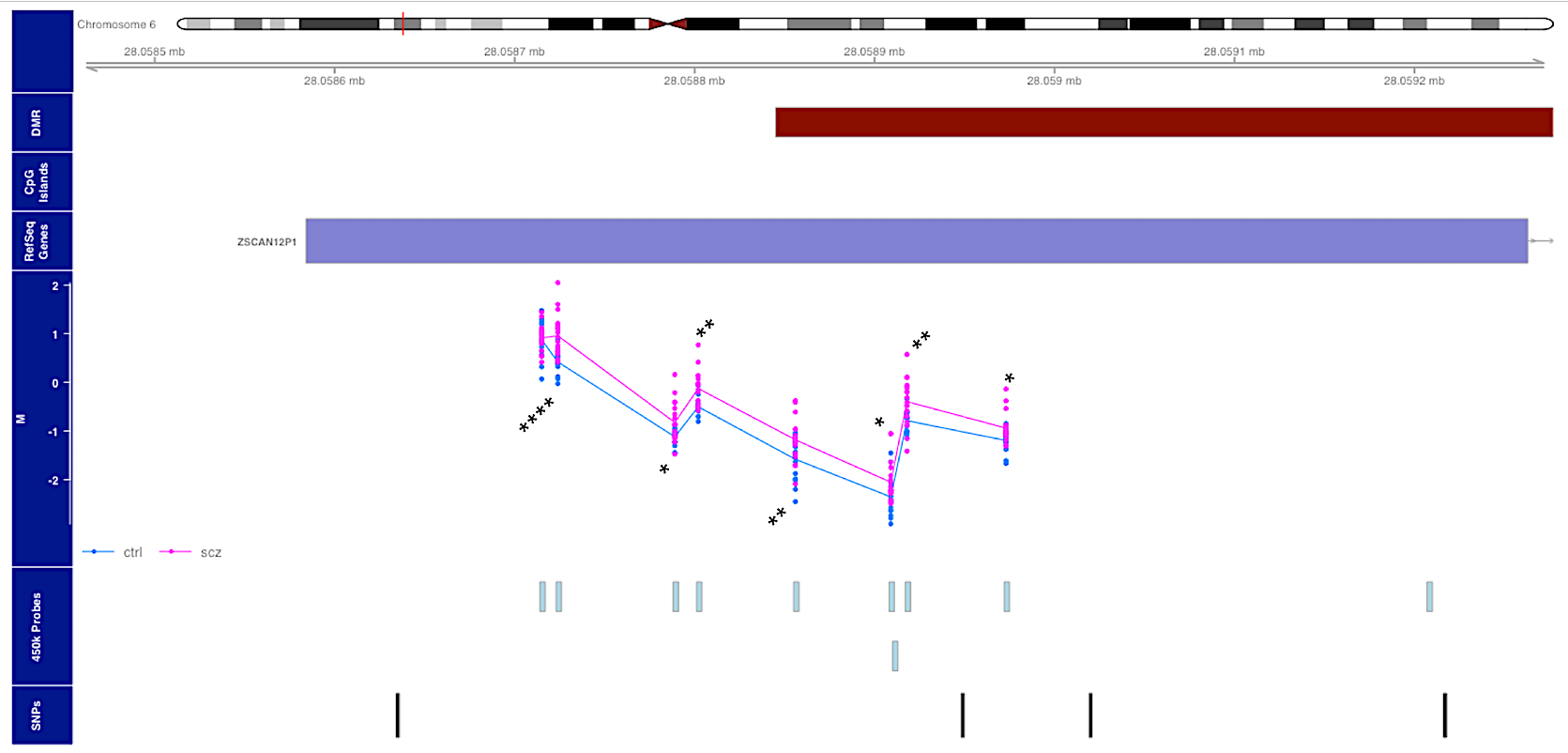

**Supplementary Figure S1** : **Methylation in patients and controls for the *DMR203* in Brain** Figure depicts methylation variation in patients with schizophrenia in the evolutionary *DMR203* (maroon, *p =* 5.064e-3 ) in the prefrontal cortex. Schizophrenia methylation (pink) compared with controls (blue) from the DBCBB dataset from Pidsley et al. Significant DMPs depicted as * (*p <*0.05), ** (*p <*0.005), and **** (*p <*0.00001)

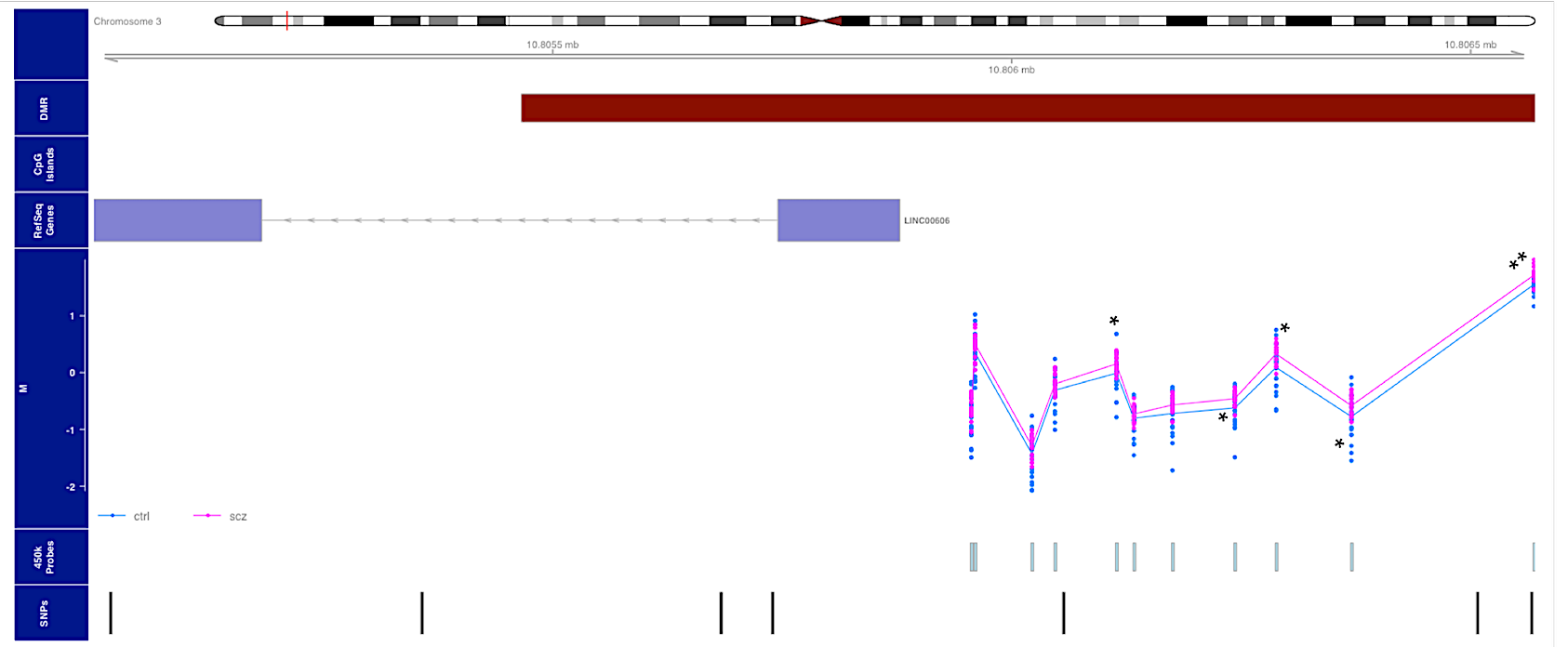

**Supplementary Figure S2** : **Methylation in patients and controls for the *DMR127* in Brain** Figure depicts methylation variation in patients with schizophrenia in the evolutionary *DMR127* (maroon, *p =* 1.078e-2) in the prefrontal cortex. Schizophrenia methylation (pink) compared with controls (blue) from the LNDBB dataset from Pidsley et al. Significant DMPs depicted as * (*p <*0.05) and ** (*p <*0.005)

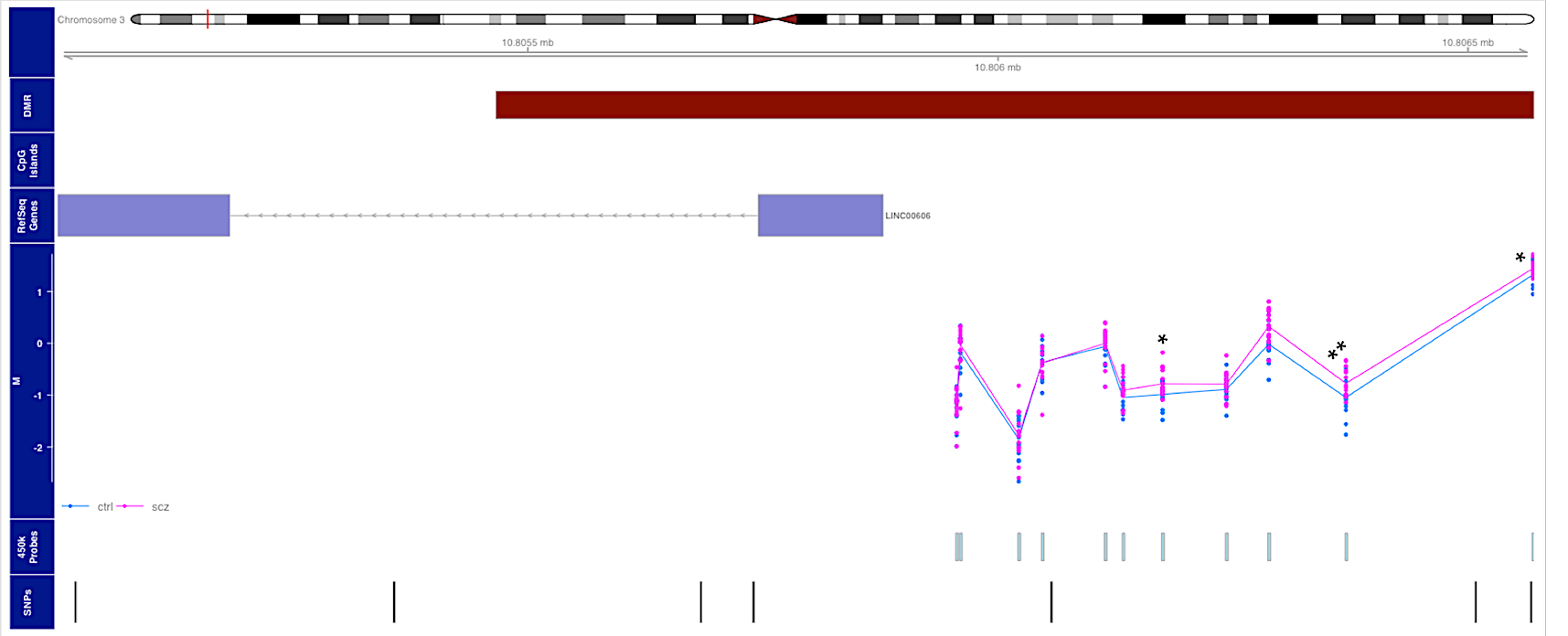

**Supplementary Figure S3** : **Methylation in patients and controls for the DMR127 in Brain** Figure depicts methylation variation in patients with schizophrenia in the evolutionary DMR127 (maroon, *p =* 0.5602) in the prefrontal cortex. Schizophrenia methylation (pink) compared with controls (blue) from the DBCBB dataset from Pidsley et al. This DMR does not reach overall significance, though it contains significant DMPs depicted as * (*p <*0.05) and ** (*p <*0.005).

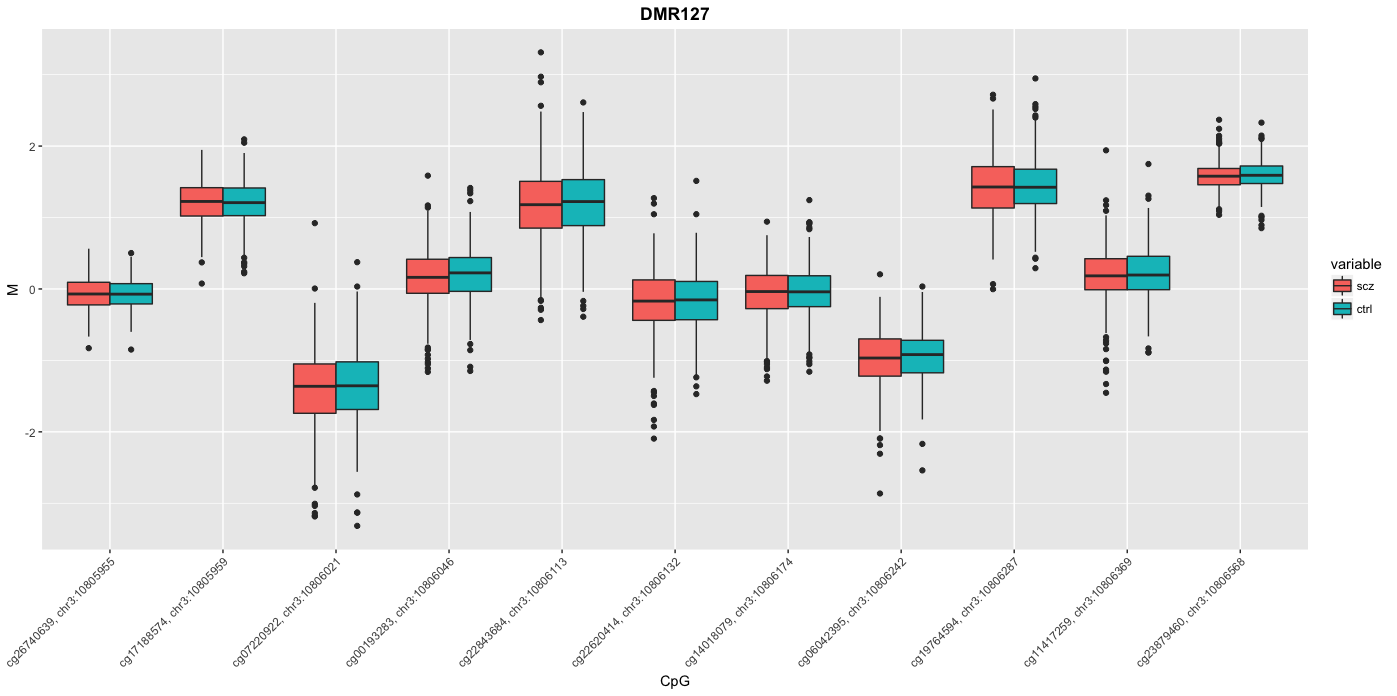

### Supplementary Figure S4: Methylation in patients and controls for the *DMR127* in Blood Figure depicts methylation variation in evolutionary *DMR127* between patients with schizophrenia (red) compared to controls (blue) in blood samples from Hannon et al. Unlike data from brain, we observe no difference in methylation from the data in blood.

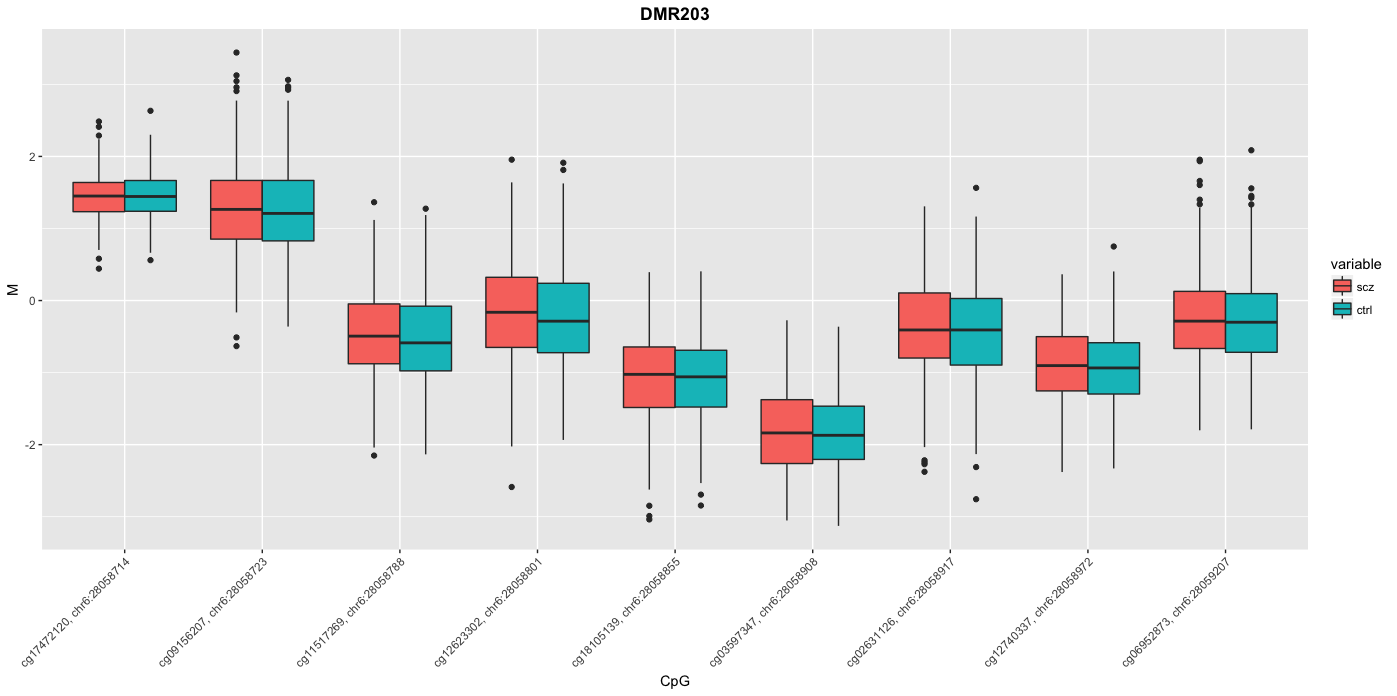

### Supplementary Figure S5: Methylation in patients and controls for the *DMR203* in Blood Figure depicts methylation variation in evolutionary *DMR203* between patients with schizophrenia (red) compared to controls (blue) in blood samples from Hannon et al. Unlike data from brain, we observe no difference in methylation from the data in blood.

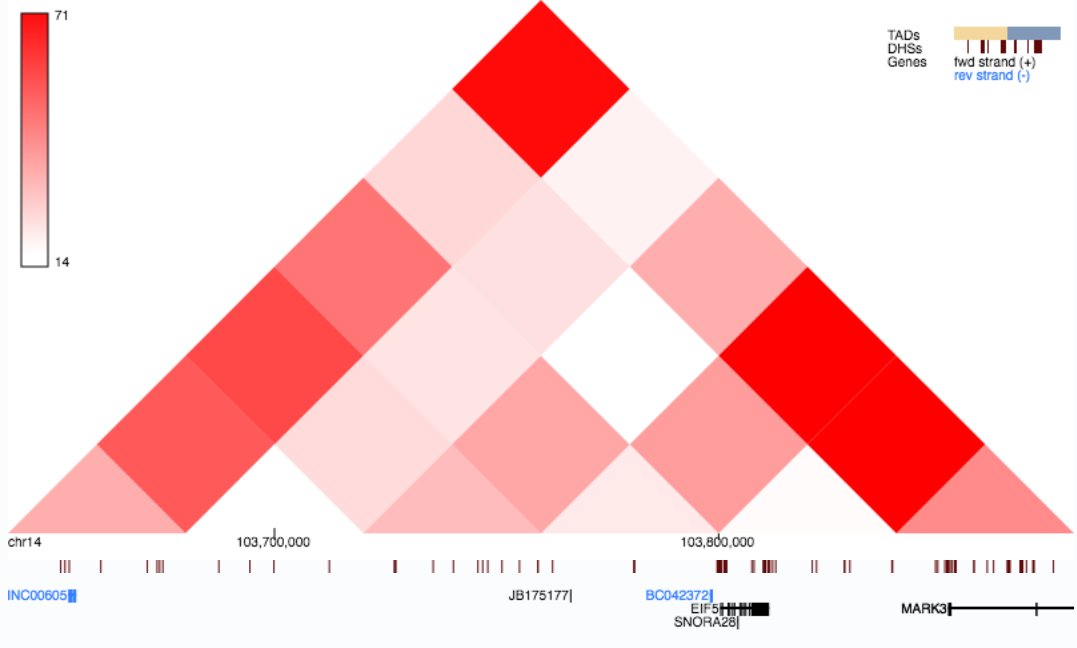

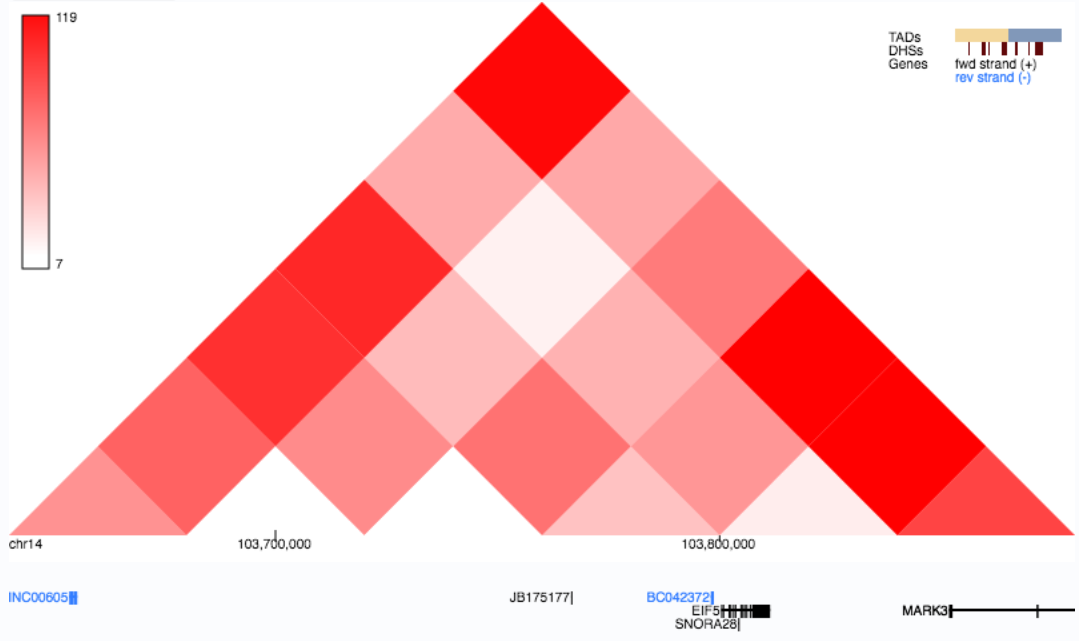

***DMR 526***

### Supplementary Figure S6: Hi-C Interaction Map for *DMR526* in Brain datasets Figure depicts a Hi-C block for evolutionary *DMR526* (blue box, hg19, chr14: 103745950-103746209) that may suggest distal regulation of MARK3*.* Top panel depicts data from hippocampus, bottom panel from dorsolateral prefrontal cortex. Hi-C data obtained from Schmitt et al, (2016).

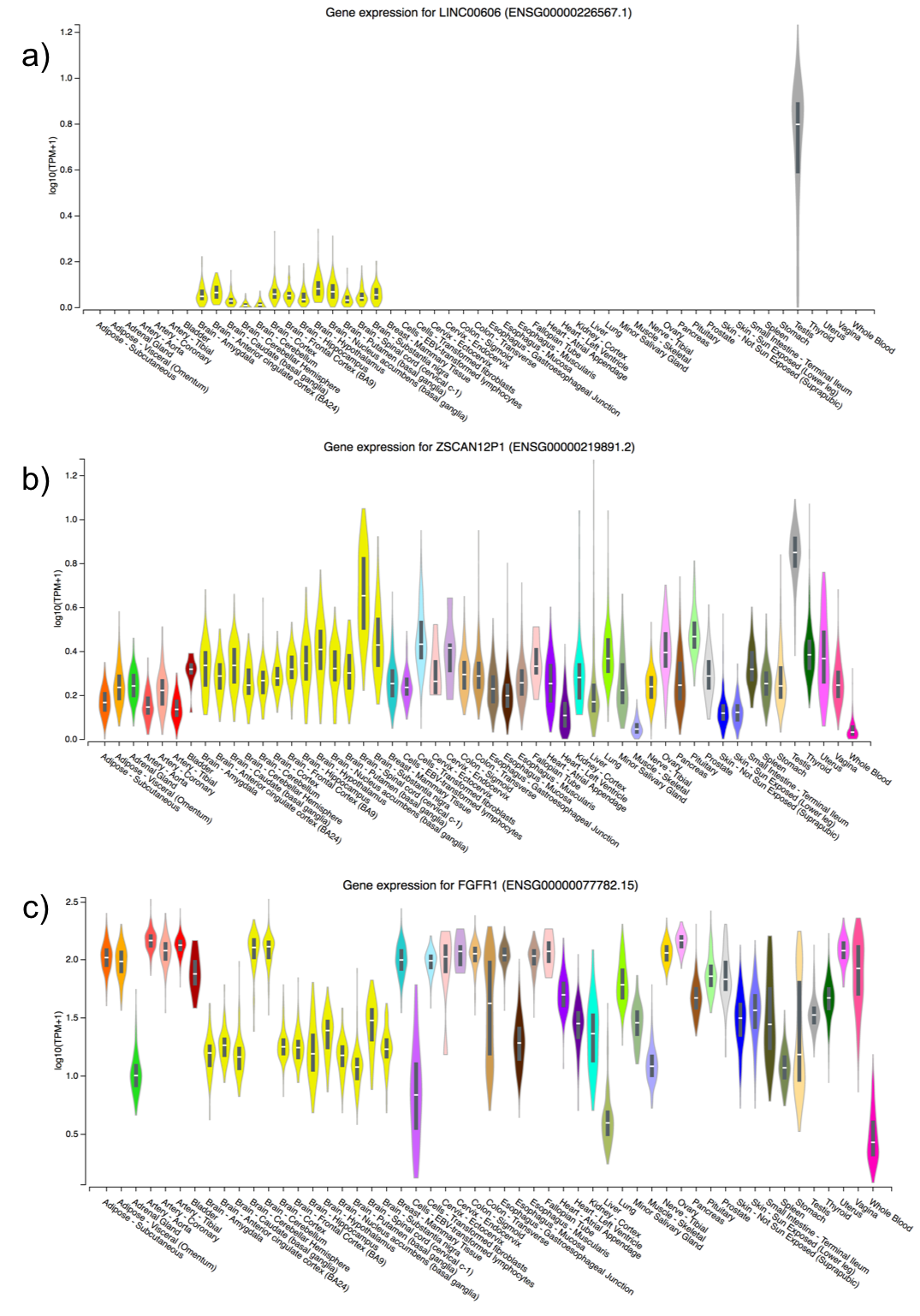

**Supplementary Figure S7**: **Global gene expression for genes overlapping evolutionary DMRs across tissues**

GTeX expression data for a) *LINC00606* b) *ZSCAN12P1* c) *FGFR1* that shows that the genes are more active in brain tissues (yellow) but not blood (magenta, extreme right side). Expression is depicted on the Y-axis as the log of Transcripts per Million (TPM). Violin plots are superimposed with box plots depicting the median , 25^th^ and 75^th^ percentile. The width of the violin plots depicts the histogram of the data.

**Supplementary Tables**

| **Sample** | **Min** | **Mean** | **Median** | **Max** |
| --- | --- | --- | --- | --- |
| Controls | 17.24 | 43.58 | 45.12 | 85.20 |
| Patients | 17.24 | 50.07 | 49.30 | 96.98 |

### Supplementary Table T1 : Age distribution in years, of samples and controls selected for analysis from Jaffe et al.

| DMR Name | SLK | Sidak |
| --- | --- | --- |
| DMR10 | 0.2052 | 1 |
| DMR127 | 0.142 | 1 |
| DMR203 | 0.7598 | 1 |
| DMR204 | 0.156 | 1 |
| DMR236 | 0.8112 | 1 |
| DMR237 | 0.1666 | 1 |
| DMR291 | 0.4057 | 1 |
| DMR526 | 0.4522 | 1 |
| DMR527 | 0.6512 | 1 |

**Supplementary Table T2 :** *Comb-P* analysis in blood data from Hannon et al. Both *Stouffer-Liptak-Kechris (Slk)* and *Sidak* correction for multiple testing are displayed with their respective *p-*values.

| DMR Name | Chromatin State  (DLPFC) | Description | Chromatin State (PBMC) | Description | Overlapping Gene |
| --- | --- | --- | --- | --- | --- |
| DMR10 | E9, E10 | Active enhancer | E11, E18 | Weak enhancer, quiescent | - |
| DMR127* | E10, E9, E10, E18 | Active enhancers, quiescent | E17 | Weak repressed polycomb | *LINC00606* |
| DMR203* | E2, E17, E18 | Flanking TSS , weak repressed polycomb, quiescent | E12, E17 | ZNF genes & repeats, weak repressive polycomb | *ZSCAN12P1* |
| DMR204 | E2, E1 | Flanking TSS, active TSS | E6, E1, E3 | Weak transcription, TSS markers | *LINC01623* |
| DMR236 | E18 | Quiescent | E5 | Strong TSS | *MAD1L1* |
| DMR237 | E6, E11 | Weak transcription + weak enhancer | E6 | Weak transcription | *MAD1L1* |
| DMR291* | E6, E11, E6 | Weak transcription + weak enhancer + weak transcription | E18 | quiescent | *FGFR1* |
| DMR526* | E15, E16 | Bivalent enhancer + repressive polycomb | E14,E16 | Bivalent TSS, repressed polycomb | - |
| DMR527 | E9, E3, E9 | Active enhancer + flanking TSS + active enhancer | E6 | Weak TSS | - |

**Supplementary Table T3 :** Epigenetic Annotation of DMRs. Data obtained by intersecting the 18-state chromatin model from Dorsolateral Prefrontal Cortex and Peripheral Blood Mononuclear Cells from Roadmap Epigenomics Project with the evolutionary DMRs, * denotes significant DMRs. Chromatin states are described in terms of emission states from ChromHMM with corresponding annotation given in the description column.
